# Transforming clinical apheresis waste into a renewable source of patient-derived CD34+ hematopoietic stem cell biobank for beta-hemoglobinopathy research and therapeutic discovery

**DOI:** 10.64898/2026.08.02.742313

**Authors:** Jun Liu, Shin-Young Park, Hirotomo Nakahara, Yembur Ahmad, Ziyang Shen, Suhayla Sarhan, Stephanie Ferrara, Vanessa Anjurthe, Kristi Georgilas, Sarah Nikiforow, Zankar Desai, Shang-Chuen Wu, Ryan P. Jajosky, Sarthak Saha, Nelya Christiansen, Kendra B. Munkacsy, Jiajia Li, Hongbo R Luo, Sophia Adamia, Sean R. Stowell, Avanish Mishra, Li Chai

**Author notes:** Co-cosrresponding authors. **CORRESPONDING AUTHORS** Jun Liu, MD, Brigham and Women’s Hospital, 75 Francis Street, Boston, MA 02115, USA, Li Chai, MD, Brigham and Women’s Hospital, 75 Francis Street, Boston, MA 02115, USA. These authors contributed equally as co-first authors (Jun Liu, Shin-Young Park, Hirotomo Nakahara, Yembur Ahmad). These authors contributed equally as co-senior authors (Avanish Mishra, Li Chai).

## Abstract

**Background aims:** The development of next-generation therapies for sickle cell disease (SCD) and beta thalassemia (beta thal), including fetal globin–inducing small molecules and gene therapy approaches, depends on patient-derived CD34+ hematopoietic stem and progenitor cells (HSPCs) for discovery and preclinical validation, but commercial vendors stock only healthy donor material and disease-specific banks hold limited inventories. Recent US Food and Drug Administration and National Institutes of Health guidance favoring human cell-based methods over animal testing underscores the value of authentic patient cells. Methods: Over 14 months we recovered, purified, and biobanked CD34+ HSPCs from clinical apheresis product waste and mobilized peripheral blood (PB) otherwise discarded after clinical procedures, using immunomagnetic selection adapted for hemoglobinopathy specimens; a microfluidic technology was evaluated separately. We quantified yield and purity for bead-selected material and cell number and viability for the microfluidic pilot; engraftment was tested in NBSGW mice. Results: Immunomagnetic selection recovered a median of 4.71 × 10^6^ CD34+ cells from just 1 to 2 mL of apheresis product waste, comparable to the 6.0 × 10^6^ cells from a 10 to 40 fold larger volume of PB waste, with similar purity across sources and diagnoses. Because apheresis product waste is far more concentrated, it reaches equivalent yields without the density-gradient steps required for PB waste, approximately halving processing time. Recovered cells engrafted NBSGW mice, confirming preserved repopulating capacity. The microfluidic pilot (two patients, 11 specimens) recovered 2.17 × 10^6^ CD34+ cells per specimen at greater than 90% viability and purity. Conclusions: A center with existing apheresis infrastructure can reproducibly recover, bank, and distribute research-grade patient CD34+ HSPCs, addressing a recognized gap in the hemoglobinopathy pipeline.

**Highlights:**

- Clinical apheresis waste is used to generate a single-center biobank of high-quality, research-grade CD34+ HSPCs from patients with sickle cell disease and beta-thalassemia.
- Concentrated apheresis waste matches large-volume PB waste in CD34+ yield and purity.
- Microfluidic enrichment recovers CD34+ cells at >90% viability and purity across 2 patients.
- Recovered CD34+ HSPCs engraft mice and form erythroid cells, preserving function.

## Introduction

Beta-hemoglobinopathies, dominated by sickle cell disease (SCD) and beta thalassemia (beta thal) but also encompassing other beta globin variants and compound heterozygotes (e.g., HbC, HbE, HbS/beta thal), together affect more than 400,000 individuals in the United States and millions worldwide [1, 2, 3]. Inherited mutations in the beta globin locus drive both diseases, and current and emerging therapies act through several mechanisms: reactivating fetal hemoglobin (HbF) to compensate for defective or absent beta hemoglobin; adding or restoring a functional beta globin gene; directly inhibiting hemoglobin S polymerization; and improving red cell survival and reducing vaso-occlusion through anti-adhesion, anti-inflammatory, and metabolic approaches [4, 5, 6, 7, 8]. Three gene therapies for SCD and/or beta thal are approved by the US Food and Drug Administration (FDA), all relying on autologous CD34+ hematopoietic stem and progenitor cells (HSPCs) collected by mobilized leukapheresis at substantial cost, resource utilization and procedural risk [9, 10]. The active development pipeline for next-generation small molecule HbF inducers, gene correction approaches, base editors, and prime editors [11, 12] depends on a steady supply of patient-sourced CD34+ HSPCs for preclinical screening, mechanism studies, and toxicology. Two recent federal policy shifts sharpen this need. In April 2025, the FDA released a roadmap to reduce, refine, or replace animal testing with human-relevant New Approach Methodologies, beginning with monoclonal antibodies and extending to other drugs [13], and the National Institutes of Health (NIH) announced that it will prioritize human-based research technologies and no longer issue funding opportunities focused exclusively on animal models [14]. As human cell-based assays displace animal studies for preclinical efficacy and toxicity testing, and as research budgets tighten, sustainable pathways to source primary patient cells become more urgent.

Small-molecule induction of HbF has become an especially active area of both preclinical and clinical development. Beyond hydroxyurea, agents now in phase 1 to 2 testing for SCD span several mechanistic classes, including DNMT1-depleting hypomethylating regimens such as oral decitabine–tetrahydrouridine, histone-modifying agents, the PRC2 (EED) inhibitor pociredir, and a new generation of cereblon-based molecular-glue degraders that eliminate the gamma-globin repressors WIZ and ZBTB7A, exemplified by the WIZ degrader ITU512 (Novartis) and the dual WIZ/ZBTB7A degrader BMS-986470 (Bristol Myers Squibb). The de-repression of gamma-globin and induction of HbF are the central pharmacodynamic endpoints for drugs in this pipeline. Healthy donor CD34+ cells from commercial vendors reproduce erythroid differentiation but fail to validate disease relevant features such as alpha/beta globin chain ratios. Therefore, the development of HbF-inducing agents depends on patient-derived CD34+ HSPCs differentiated along the erythroid lineage and/or in humanized xenotransplantation models. The molecular-glue degrader of WIZ, for example, was validated in patient CD34+ HSPC-derived erythroid cultures [15]. As we developed our own HbF-inducing small molecule degrader in-house [16, 17], limited access to patient cells repeatedly slowed our pace. The same constraint applies to other candidates in early development [16]. A renewable supply of disease-relevant patient CD34+ cells is therefore a prerequisite for advancing these promising classes of therapeutics.

A recent Cure Sickle Cell Initiative Apheresis Working Group review by Tanhehco and colleagues catalogued the clinical challenges of hematopoietic stem cell mobilization in SCD: patients with SCD carry a steady state PB CD34+ count well below that of healthy donors (median 7.3 cells per microliter), use of granulocyte colony-stimulating factor (G-CSF) carries a risk of severe vaso-occlusive crisis, and gene therapy manufacturing requires 15 × 10^6^ CD34+ cells per kilogram as starting material [18]. Mobilizing a patient with SCD for a clinical gene therapy collection takes considerable preparation and carries real procedural risk, and because gene therapy has not yet expanded widely only a small subset of patients undergo these collections. Every collection is therefore valuable, and full use of each one, including banking of otherwise discarded waste material for research, recovers material that would otherwise be lost.

Three sources currently supply patient-sourced HSPCs. The first is the National Heart, Lung, and Blood Institute (NHLBI) Cure Sickle Cell Initiative Sickle Cell Hematopoietic Stem Cell Bank (BioLINCC accession HLB02662222a), which provides plerixafor-mobilized CD34+ and CD34-negative cells from a small set of SCD volunteers [19]. The second is bone marrow biopsy, which is rarely performed for this purpose and yields only small numbers of cells. The third is apheresis waste material obtained ad hoc by individual investigators: because only a subset of US academic medical centers operate apheresis units that perform SCD gene-therapy CD34+ collections, interested laboratories must identify such a center, negotiate the logistics of transferring waste blood with the apheresis practice, and then carry out their own CD34+ selection. Each source is useful but insufficient for current demand, and rarely scalable to industry partnerships.

Here we describe a fourth source: a single-center platform that systematically captures clinical apheresis and peripheral blood (PB) waste from routine apheresis procedures, and recovers, characterizes, and biobanks CD34+ HSPCs and matched CD34-negative mononuclear cells. The platform builds on the Brigham and Women’s Hospital (BWH) Apheresis Research Core, which supports more than thirty clinical trials in oncology, autoimmune disease, and infectious disease and is capable of providing fresh cells from apheresis waste for research [20]. We extend this infrastructure to hemoglobinopathies and, in collaboration with the Li Chai and Avanish Mishra laboratories, report measured yields, purity, viability, and functional validation from 14 months of operation, together with a research microfluidic enrichment pilot. We describe how other institutions could replicate the technical workflow.

## Methods

### Patient consent and Institutional Review Board framework

A master research protocol approved by the Mass General Brigham Human Research Committee governs the retrieval of waste material. Consent for general research use is already embedded in the standard clinical apheresis consent form, which permits a defined set of general research uses (*in vitro* erythroid, myeloid, and lymphoid differentiation; drug efficacy and toxicity testing; *in vivo* xenotransplantation; disease mechanism investigations; testing of new CD34 isolation devices; and cryopreservation and biobanking). Special use cases that fall outside routine clinical research require an additional research consent which can be obtained separately through the Apheresis Research Core, covering human leukocyte antigen (HLA) typing, generation of induced pluripotent stem cells (iPSCs), broad molecular profiling (whole genome sequencing, RNA sequencing, and proteomics), genetic modification (CRISPR or base editing), microbiome and infectious disease studies, and sharing with external academic or industry collaborators.

### Apheresis Research Core to local lab operations

We collect material from adult patients with SCD or beta thal who undergo clinically indicated mobilized apheresis procedures at the BWH Kraft Family Blood Donor Center. Two specimen streams enter the workflow: (i) “apheresis product waste”, typically 1 to 2 mL of residual cell-rich material from the leukopak bulb after disconnection, and (ii) “unused mobilized PB waste”, which are typically 30 to 70 mL remaining in tubing or unused diversion. PB specimens are sent to the Chai laboratory within two hours of collection; apheresis product waste is kept at 4 °C in ethylenediaminetetraacetic acid (EDTA) tubes at the cell manufacturing facility and sent to the Chai laboratory the following morning.

Operational governance of these pathways is handled by the Apheresis Research Core. Core director or quality/compliance specialist reviews requests before any material is released. Requests follow a standard requisition process in which the investigator submits a requisition form, and the Apheresis Research Core then notifies the clinical team and apheresis nurses to save the relevant specimens, and specimens are logged at retrieval so that each can be traced and its disposition documented in accordance with AABB standards. The product distribution scheme and the full consent, compliance, and governance workflow are detailed in a separate manuscript.

### Patient mobilization and recorded clinical variables

Patients are mobilized per institutional practice: adults with SCD receive plerixafor or motixafortide, and adults with beta thal receive G-CSF with either plerixafor or motixafortide. Under the master Institutional Review Board (IRB) protocol we record basic demographic and clinical variables that are important for interpreting the functional *in vitro* and *in vivo* results, including sex, age range, and hydroxyurea exposure and timeline.

### Mononuclear cell isolation and red blood cell lysis

PB waste undergoes density gradient separation: we dilute the specimen in phosphate-buffered saline containing 2% fetal bovine serum, layer it over Lymphoprep (StemCell Technologies), and centrifuge at 800 g for 30 minutes at room temperature to recover the buffy coat. Apheresis product waste, which is already cell-rich, bypasses density gradient separation and proceeds directly to red blood cell lysis per the EasySep protocol. We then wash the recovered mononuclear cells once in selection buffer, count them on an automated hemocytometer, and assess viability by trypan blue exclusion.

### CD34+ immunomagnetic selection

We enrich CD34+ HSPCs using the EasySep Human CD34 Positive Selection Kit II, catalog 17856, or the higher purity catalog 17879 (StemCell Technologies), per the manufacturer protocol [21], with two adaptations that we developed for this cohort: (i) four to five sequential incubation and wash cycles per selection (versus the three cycles in the standard protocol) to compensate for the higher debris and granulocyte content of apheresis waste, and (ii) batched processing of large mononuclear cell fractions in 5 × 10^8^ cell aliquots when the total mononuclear cell count exceeds 10^9^ cells.

### Cryopreservation and biobanking

We resuspend CD34+ cells at 1 to 5 × 10^6^ cells per mL in CryoStor CS10 (BioLife Solutions) [22] or 10% dimethyl sulfoxide (DMSO) in heat-inactivated fetal bovine serum, freeze them at a controlled rate to minus 80 °C, and transfer them to the vapor phase of liquid nitrogen for long-term storage. We bank CD34-negative mononuclear cell fractions in parallel in CryoStor CS10 and in 10% DMSO in fetal bovine serum to support immunology and iPSC reprogramming studies [23].

### Research microfluidic enrichment

As a pilot, we ran an alternative enrichment workflow based on a research microfluidic technology currently in development by an internal collaborator, on 11 specimens from two patients with SCD. This approach enriches CD34+ cells directly from PB or apheresis waste without red blood cell lysis. For the recovered CD34+ cells, we assessed cell yield and viability as for bead-selected material; a detailed characterization of this technology will be reported separately.

### Quality assessment

We measure post-selection cell counts on an automated hemocytometer and assess CD34 purity by in-house flow cytometry, staining cells with a conjugated anti-CD34 antibody (BD Biosciences) and acquiring data on a Cytek cytometer alongside an isotype control.

### Functional validation

#### In vitro erythroid differentiation

CD34+ HSPCs from a patient with SCD and a patient with beta thal were differentiated along the erythroid lineage using a three-phase liquid culture protocol adapted from published methods [17]. Cells were recovered in erythroid differentiation medium (Iscove’s Modified Dulbecco’s Medium supplemented with holo-human transferrin, recombinant human insulin, heparin, inactivated plasma, erythropoietin, and L-glutamine); the medium was supplemented with hydrocortisone, stem cell factor, and interleukin-3 for the first phase (7 days), with stem cell factor alone for the second phase (4 days), and without additional supplements for the final phase (5 to 7 days). Parallel cultures were treated with an investigational HbF-inducing compound (Compound X; 0.05 uM) or vehicle control (DMSO), and erythroid maturation was monitored over time by flow cytometry for the CD235a+ (glycophorin A) population.

#### Human CD34+ cell xenotransplantation

Reconstitution capacity of recovered HSPCs was assessed by xenotransplantation following the protocol of Liu and colleagues [17]. NBSGW mice (NOD.Cg-Kit^W-41J Tyr^+ Prkdc^scid Il2rg^tm1Wjl/ThomJ; Jackson Laboratory stock 026622), 6 to 8 weeks of age and maintained as homozygotes in a pathogen-free facility, were conditioned with low-dose busulfan (10 mg/kg) and transplanted by intravenous or retro-orbital injection with 1 × 10^5^ immunomagnetically selected CD34+ HSPCs pooled from the PB and apheresis waste of a single patient with SCD. Human hematopoietic reconstitution was monitored longitudinally by flow cytometric analysis of PB, measuring human CD45+ (hCD45+) leukocyte chimerism at 4, 8, and 12 weeks after transplantation on a Attune NxT cytometer (Thermo Fisher). All animal procedures were approved by the Institutional Animal Care and Use Committee (IACUC).

### Statistical analysis

We report medians and ranges for cell yield and purity. Two selections that were determined to have counting errors were excluded from all yield and purity analyses and from all figures; all other measured selections were retained. Because group sizes were small, we compared groups using the two-sided Mann-Whitney U test (exact where feasible) using SciPy, and considered p < 0.05 as significant.

## Results

### Platform overview

We assembled an integrated workflow that funnels two categories of clinical waste material, mobilized PB and apheresis product waste, through density gradient mononuclear cell isolation for PB waste or direct red blood cell lysis for apheresis waste, followed by immunomagnetic CD34+ enrichment, and parallel banking of viable CD34+ HSPCs and CD34-negative mononuclear cells. A research microfluidic technology was evaluated separately as an alternative enrichment route. The resulting biobank supports drug discovery, gene therapy preclinical work, drug toxicity testing, and sequencing studies (**Figure 1**).

**Figure 1.**
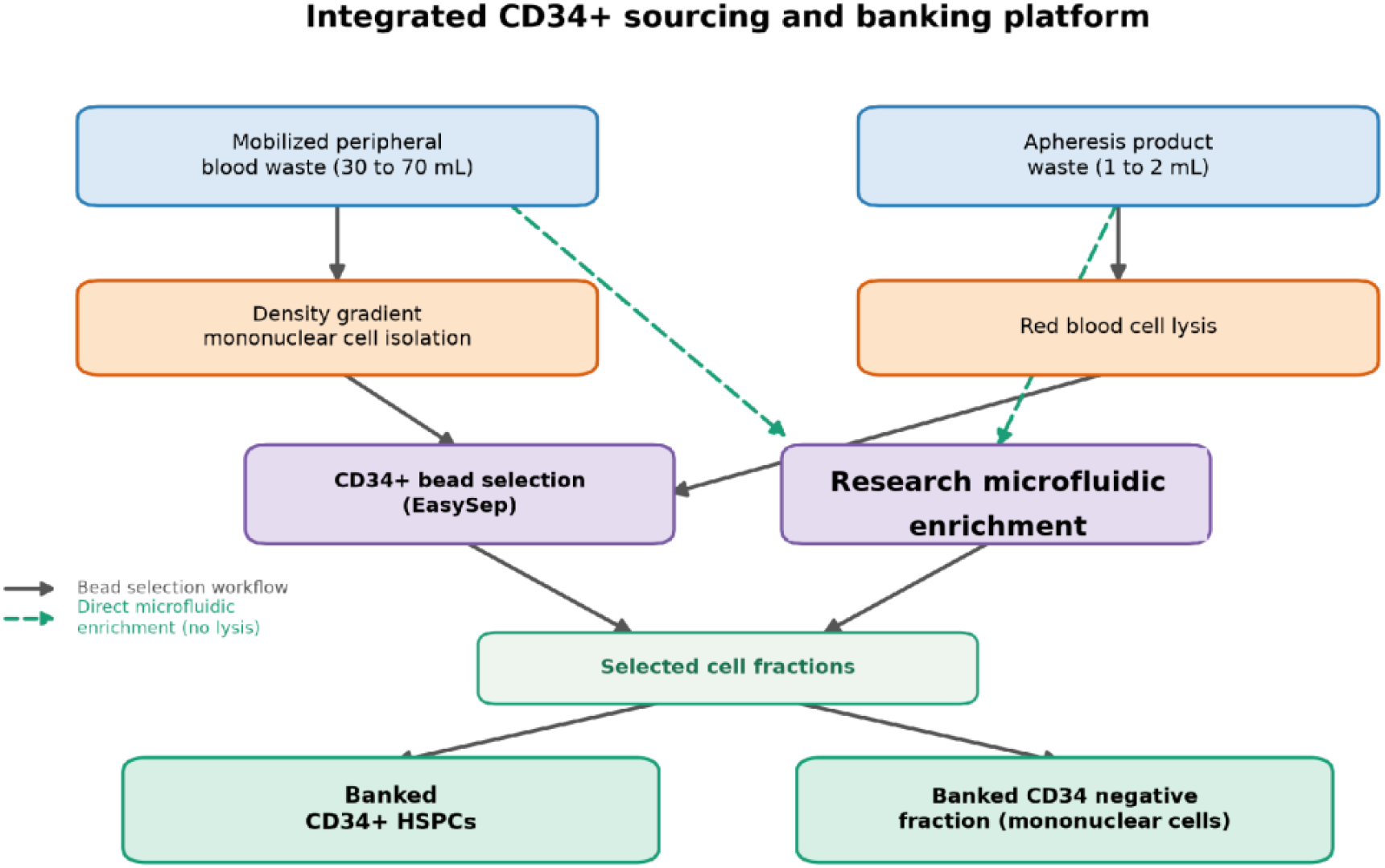
Integrated CD34+ sourcing and banking platform. Patients consented under a master IRB framework contribute two categories of clinical waste through the BWH Apheresis Research Core. The workflow processes PB waste by density gradient mononuclear cell isolation and apheresis waste by direct red blood cell lysis, followed by CD34+ immunomagnetic selection or research microfluidic enrichment, then distributes parallel banked products for downstream applications.

### Efficient recovery of patient-derived CD34+ HSPCs

Between January 2025 and March 2026, we performed 30 CD34+ cell selections from 10 unique patients with SCD or beta thal (4 with SCD, 6 with beta thal) alongside 6 healthy donor controls used to monitor workflow stability. Two selections that were determined to have counting errors were excluded from all yield and purity analyses. Apheresis product waste specimens, despite volumes of only 0.8 to 5.5 mL (median 1.3 mL), delivered a median of 4.71 × 10^6^ CD34+ cells per selection (range 1.3 to 11.2 × 10^6^, n = 19; Figure 2A). Mobilized PB waste of 12.5 to 47.5 mL yielded a median of 6.0 × 10^6^ CD34+ cells per selection (range 4.26 to 9.0 × 10^6^, n = 4). Despite requiring roughly 10 to 40-fold less starting volume, apheresis product waste therefore matched PB waste in CD34+ yield (median 4.71 versus 6.0 × 10^6^; p = 0.16), a direct consequence of its far higher CD34+ concentration (1,000 to 6,000 versus 50 to 200 cells per µL). Recovery was also comparable between SCD and beta thal specimens (median 4.10 × 10^6^ versus 5.00 × 10^6^ CD34+ cells; p = 0.64), indicating that the platform performs robustly across both disease types and specimen sources.

**Figure 2.**
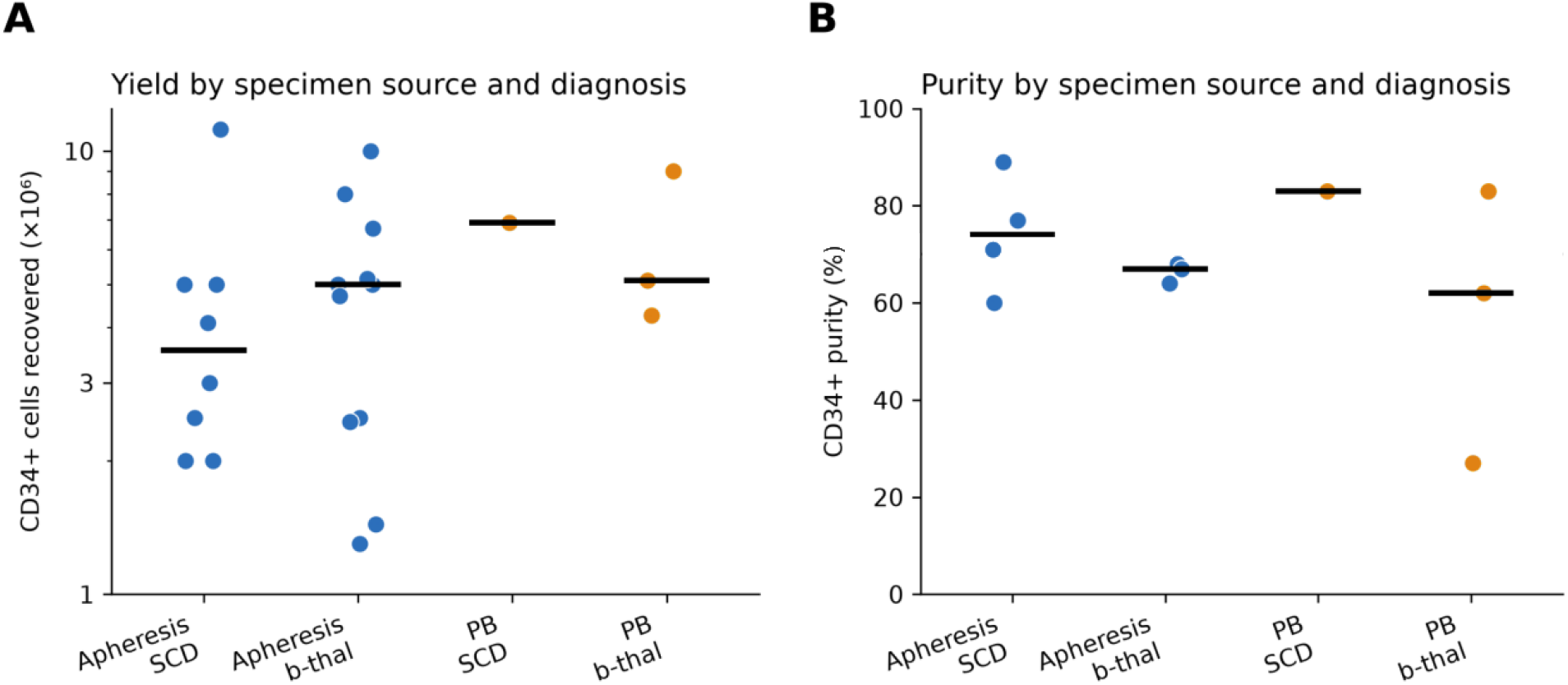
CD34+ HSPC yield and purity by bead-based (EasySep) immunomagnetic selection. (A) CD34+ HSPC yield by specimen source and diagnosis. Raw CD34+ cells recovered per selection by bead-based (EasySep) immunomagnetic selection, shown on a log scale for apheresis and peripheral blood (PB) waste from patients with SCD or beta thal; horizontal bars are group medians. Two selections flagged as counting errors were excluded. n = 23 patient selections from 10 unique patients. (B) CD34+ purity by specimen source and diagnosis for bead-based immunomagnetic selection. In-house flow cytometry purity for apheresis and PB waste from patients with SCD or beta thal; horizontal bars are group medians. n = 11 selections with a purity measurement.

**Figure 3.**
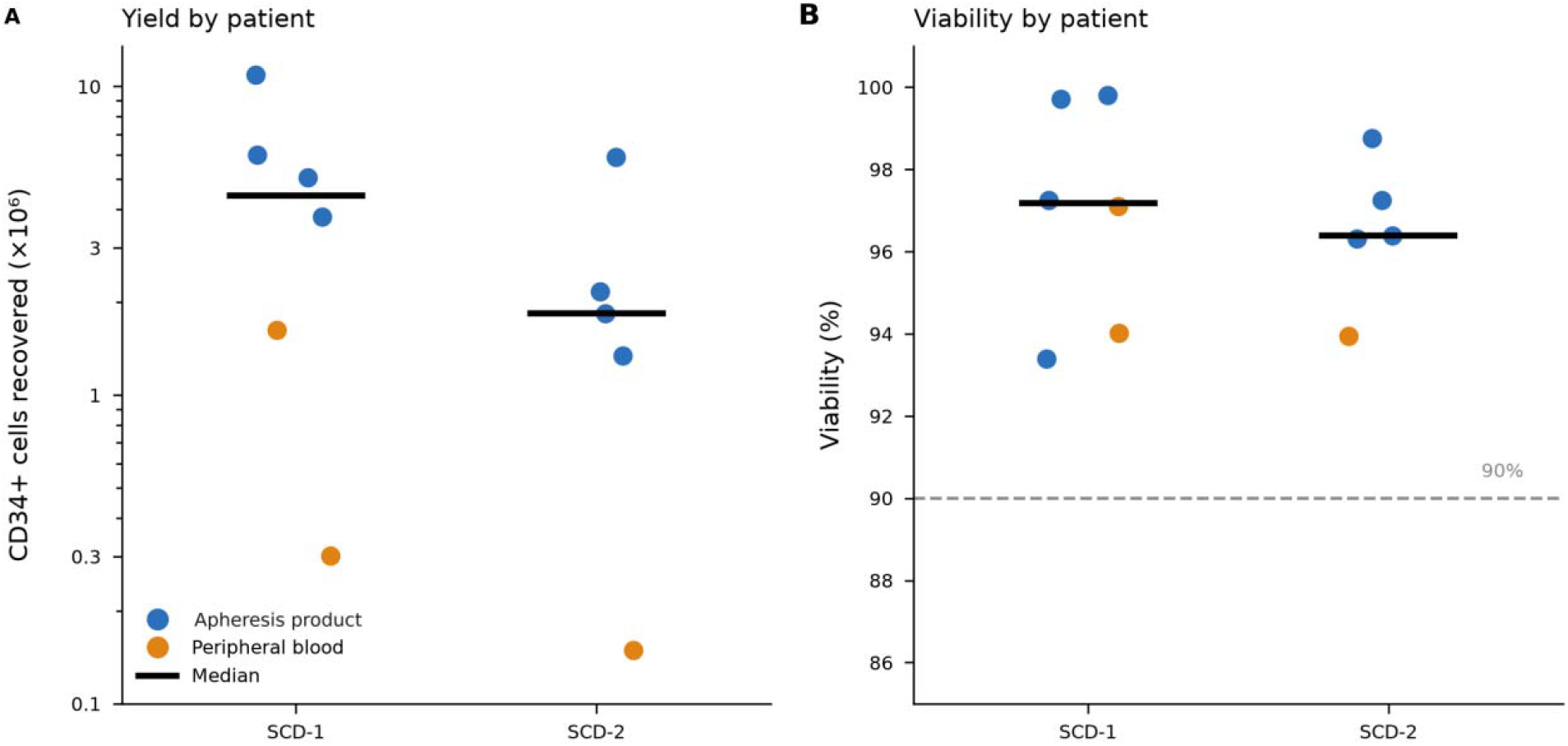
CD34+ HSPC recovery by a research microfluidic technology. (A) CD34+ cells recovered per specimen (log scale) for two patients with SCD (SCD-1, n = 6; SCD-2, n = 5); points are individual specimens colored by input material (apheresis product or PB), horizontal bars are patient medians. (B) Post enrichment viability by trypan blue exclusion for the same specimens; dashed line marks 90%. n = 11 specimens from 2 unique SCD patients.

### Recovered HSPCs retain moderate purity

CD34+ purity by in-house flow cytometry varied modestly by source and diagnosis (**Figure 2B**): apheresis SCD specimens reached a median of 74% (range 60 to 89%, n = 4), apheresis beta thal specimens a median of 67% (range 64 to 68%, n = 3), and PB beta thal specimens a median of 62% (range 27 to 83%, n = 3); the single PB SCD specimen with a purity measurement reached 83%. Purity did not differ significantly between apheresis and PB waste (p = 0.93) or between diagnoses (p = 0.25). Purity was modestly lower than that typically observed in healthy donor collections, consistent with the higher background of activated cells and debris in hemoglobinopathy specimens; these differences did not compromise downstream applications. Cells were cryopreserved using standardized clinical protocols and maintained as a renewable biobank for subsequent experimental use.

### Functional validation of recovered HSPCs

Selected SCD patient CD34+ HSPCs were assessed for *in vivo* engraftment competence by intravenous or retro-orbital transplantation of 1 × 10^5^ immunomagnetically selected CD34+ HSPCs, pooled from the PB and apheresis waste of a single SCD patient, into NBSGW mice (n = 8 recipient mice; 5 male, 3 female). Human CD45+ chimerism was measurable in all recipients and rose from a mean of 3.0% at 4 weeks to 8.1% at 8 weeks, with 6.3% maintained at 12 weeks; individual recipients reached up to 14.2% at 8 weeks and 10.5% at 12 weeks (Figure 4).

**Figure 4.**
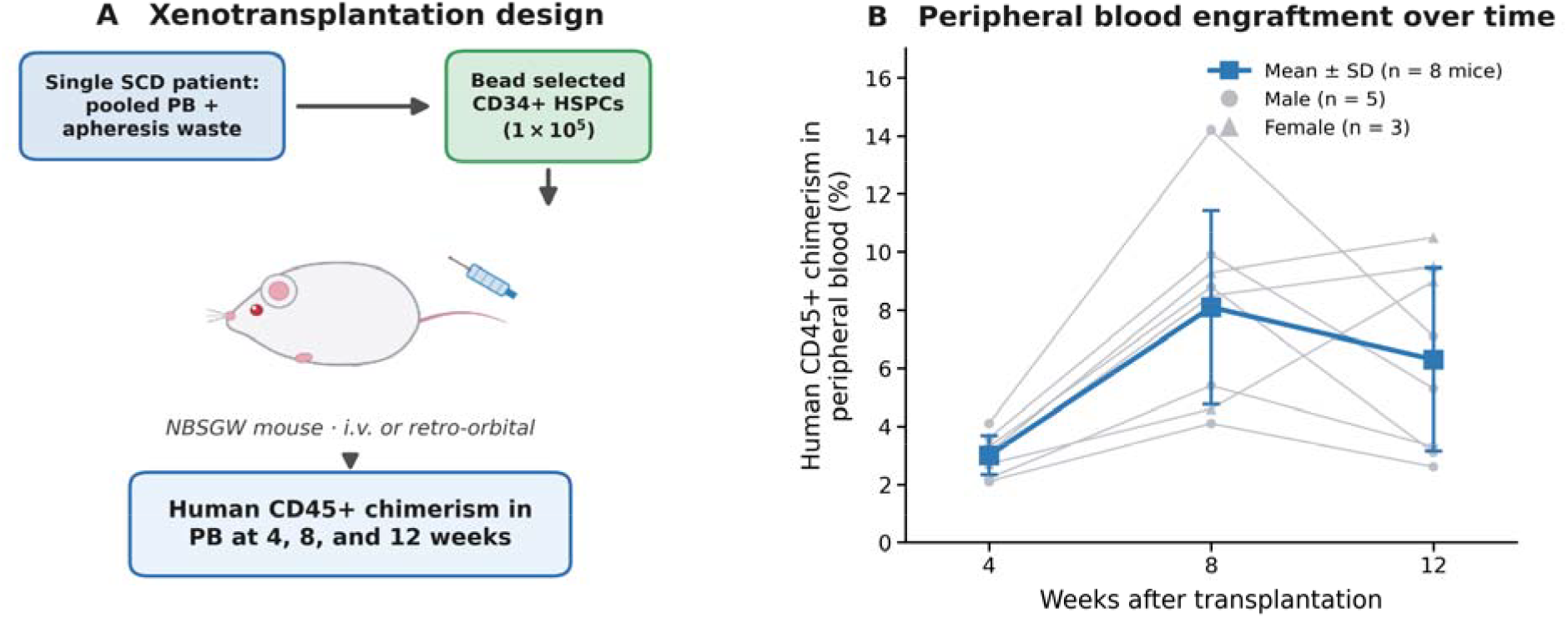
SCD patient CD34+ HSPC engraftment in NBSGW mice, measured in PB. (A) Xenotransplantation design: 1 × 10^5^ immunomagnetically selected CD34+ HSPCs pooled from the PB and apheresis waste of a single SCD patient were injected (i.v./retro-orbital) into NBSGW mice, and human CD45+ chimerism was sampled from PB at 4, 8, and 12 weeks following the protocol of Liu, Shen, Park and colleagues [17]. (B) Human CD45+ chimerism in PB over time (n = 8 recipient mice; 5 male, 3 female). Thin grey lines are individual recipients (circles, male; triangles, female); the bold blue line is the group mean with error bars showing standard deviation.

Because chimerism was measured in circulating PB rather than at a single bone marrow endpoint, these results indicate that CD34+ HSPCs recovered from SCD apheresis waste retain repopulating capacity detectable in the circulation through 12 weeks. Recovered CD34+ HSPCs also underwent *in vitro* erythroid differentiation: cells from a patient with SCD and from a patient with beta thal, treated with the HbF-inducing lead Compound X, generated CD235a+ erythroid populations comparable to controls (Figure 5), confirming preserved erythroid differentiation capacity after recovery and cryopreservation.

**Figure 5.**
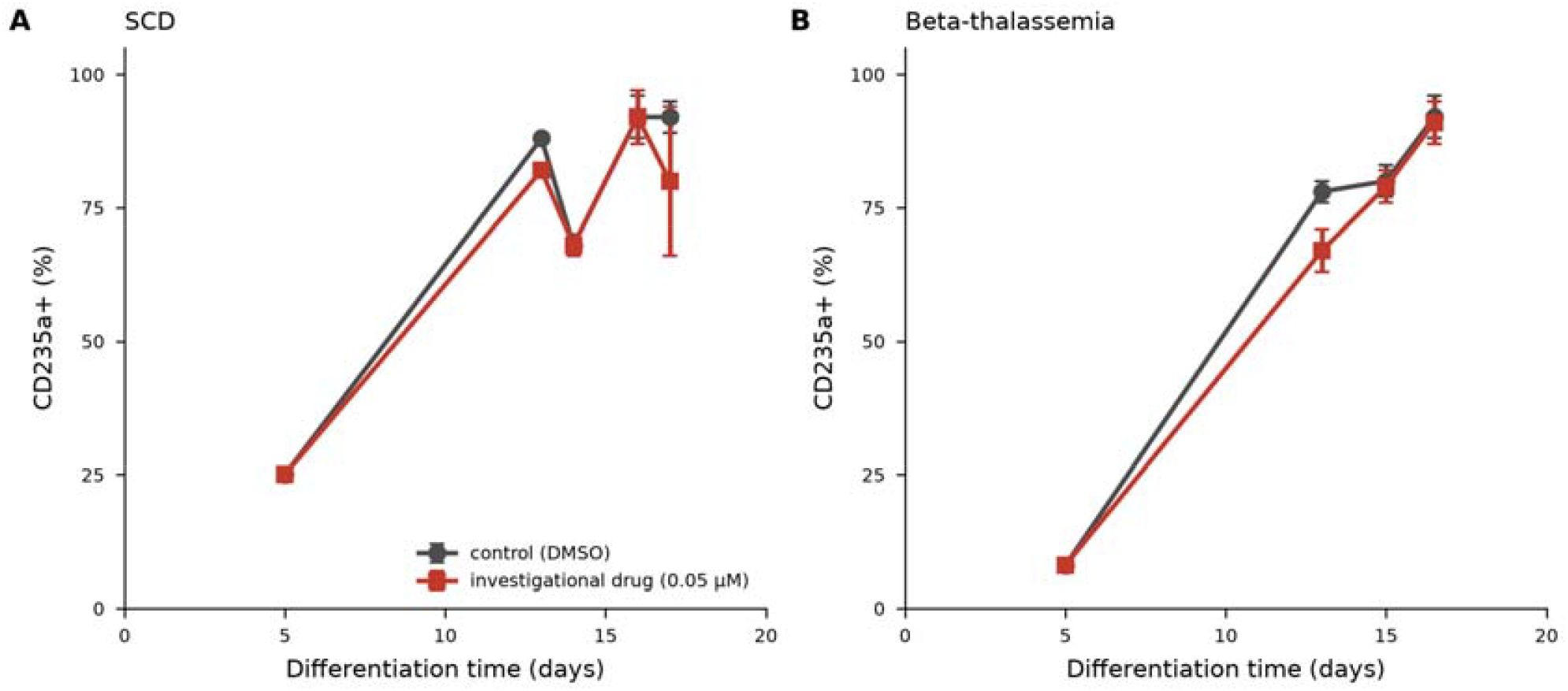
*In vitro* erythroid differentiation of recovered patient CD34+ HSPCs. CD34+ HSPCs from a patient with SCD and a patient with beta thal were differentiated along the erythroid lineage in the presence of an investigational HbF-inducing drug (0.05 µM) or vehicle control (DMSO), and the CD235a+ erythroid fraction was monitored over time by flow cytometry. (A) SCD: percentage of CD235a+ cells versus days of differentiation for control and drug-treated cultures. (B) Beta thal: the corresponding CD235a+ time course. In both diagnoses the CD235a+ fraction rises from approximately 10 to 25% early in differentiation to roughly 90% by days 15 to 17, and the drug-treated and control trajectories overlap closely, indicating that erythroid differentiation of recovered CD34+ HSPCs is preserved under drug treatment. Values are group means (error bars show standard deviation where available).

### Microfluidic enrichment recovers CD34+ HSPCs at high viability and purity

As an alternative enrichment route, a research microfluidic technology was applied to 11 specimens from two additional patients with SCD. This approach recovered a median of 2.17 × 10^6^ CD34+ cells per specimen (range 0.15 to 10.91 × 10^6^) at 96.4% purity (range 93.9% to 98.3%), with apheresis residual specimens yielding more than direct PB draws (median 4.42 versus 0.30 × 10^6^ CD34+ cells). Median volume of processed apheresis residual specimens was 1 mL and 5 mL for direct PB draws. Recovered cells showed a median viability of 97.1% (range 93.4 to 99.8%; Figure 3). Although evaluated in a limited pilot, these results indicate that microfluidic enrichment recovers viable CD34+ HSPCs directly from PB or apheresis waste and merits fuller characterization.

## Discussion

The rapid expansion of gene-editing technologies, targeted therapeutics, and cell-based treatments has transformed the therapeutic landscape for SCD and beta thal. Regardless of modality, nearly all preclinical pipelines have a common requirement: access to functional patient-derived HSPCs. Although numerous studies have identified promising targets for HbF induction and gene correction [4, 15, 16, 17, 24], the availability of authentic patient-derived CD34+ HSPCs has become an increasingly important rate-limiting step for translating these discoveries into clinically relevant therapies. This constraint is becoming more pressing as the FDA and NIH guidance shifts preclinical efficacy and toxicity testing away from animal models toward human cell-based New Approach Methodologies [13, 14]. In this study, we show that residual clinical apheresis material, routinely discarded after stem cell collection, is a robust and renewable source of functional patient-derived HSPCs that can be systematically recovered, cryopreserved, and deployed for translational research.

Relatively small volumes of apheresis product waste material consistently yielded millions of viable CD34+ cells suitable for downstream applications. Notably, the small-volume, highly concentrated apheresis product waste (1 to 2 mL) matched the larger-volume PB waste (12.5 to 47.5 mL) in both CD34+ yield and purity despite containing 10 to 40-fold less starting material. This concentration advantage has a practical operational consequence: PB waste must first undergo density-gradient (Ficoll) centrifugation and mononuclear cell enrichment before selection, which adds roughly 2 to 3 hours and approximately doubles hands-on processing time relative to apheresis product waste, which is enriched directly without a mononuclear cell isolation step. Apheresis product waste is therefore the preferred input where available. For centers with limited bandwidth for waste material processing, apheresis product waste (readily available since an apheresis specimen is always taken for CD34 enumeration post-collection), thus offers a more cost-effective biobank starting material compared to PB waste.

Importantly, recovered cells retained hematopoietic function after cryopreservation, as shown by engraftment in NBSGW mice and by *in vitro* erythroid differentiation of both SCD and beta thal specimens. These observations indicate that this clinical waste material can support studies that require functional hematopoietic stem cells. Because these specimens originate directly from patients undergoing clinically indicated collection, they preserve disease-specific biological characteristics that cannot be fully reproduced using healthy donor cells or immortalized erythroid lines. A research microfluidic technology offers a complementary enrichment route that operates directly on PB or apheresis waste and recovered viable CD34+ HSPCs in a pilot across two patients; its fuller functional characterization is a priority for future work.

Patient-derived CD34+ HSPCs recovered through this platform have supported 5 studies of HbF-inducing therapeutics, xenotransplantation, erythroid differentiation, mechanistic investigations of hematopoietic stem cell biology, and novel device development. Research groups include both internal (Harvard) and external collaborators. Parallel banking of matched CD34-negative mononuclear cells further expands the utility of this approach for immunologic, genomic, and functional studies. Collectively, these findings establish that residual clinical leukapheresis waste material is a robust and renewable source of functional patient-derived HSPCs, providing an enabling resource for translational studies aimed at developing next-generation therapies for SCD and beta thal.

## Limitations

This study has several limitations. First, the current cohort represents a single-center experience with a relatively modest number of patients (n = 11); larger multicenter studies will be necessary to evaluate reproducibility across diverse patient populations and collection practices. Second, CD34+ purity from patient apheresis waste falls below that of healthy donor controls and may underestimate functional HSPC content; matched single-cell analyses will further define the functional HSPC fraction in our cohort [25]. Finally, functional validation to date has used immunomagnetically selected cells; patient CD34+ HSPCs isolated by the research microfluidic technology have not yet been carried through *in vitro* erythroid differentiation or *in vivo* engraftment, and confirming their functional equivalence is a priority for future work. A further practical limitation is that cryopreservation and biobanking currently depend on capacity within the Chai laboratory, which is resource-intensive and costly; distributing this step or establishing dedicated shared biobanking infrastructure would improve scalability and cost-efficiency.

## Acknowledgements

We thank the patients who consented to research use of clinical waste material; the Apheresis Research Core staff at the Kraft Family Blood Donor Center; the Brigham and Women’s Hospital Transfusion Medicine team; the Dana Farber Cancer Institute Cell Manipulation Core Facility (CMCF) team; and the Mass General Brigham Human Research Committee. We also thank Junyan Zhang, Danielle Tenen, Po-Shen Chen for technical support, and the laboratories of Daniel Bauer, Daniel Tenen, John Manis, David Justus, and Avanish Mishra for ongoing collaboration.

## Author Contributions

Jun Liu and Li Chai conceptualized the project, with significant input from Hirotomo Nakahara, Yembur Ahmad, and Sean Stowell. Avanish Mishra and Sarthak Saha developed and validated the microfluidic isolation method and provided cellular data. Shin-Young Park and Jiajia Li performed *in vivo* validation. Shang-Chuen Wu and Ryan P. Jajosky provided crucial support for flow cytometry validation of cell properties. Jun Liu created the manuscript draft, with significant input from Li Chai, Avanish Mishra, and Yembur Ahmad. All other authors significantly contributed to the development and validation of the sample requisition and processing pipeline.

## Data Availability Statement

The data that support the findings of this study are available from the corresponding author upon reasonable request.

## Ethics Approval and Consent

All work was conducted under Mass General Brigham Human Research Committee IRB protocol 2025P000309; all patients provided written informed consent for research use of clinical waste material. Animal studies were approved under IACUC protocol 102954.

